# Phosphorylation of light-harvesting complex II controls excitation energy spillover between photosystems

**DOI:** 10.1101/2022.07.10.499486

**Authors:** Ryutaro Tokutsu, Eunchul Kim, Seiji Akimoto, Makio Yokono, Konomi Fujimura-Kamada, Norikazu Ohnishi, Yoshifumi Ueno, Jun Minagawa

## Abstract

Land plants and microalgae convert solar energy into electrochemical energy by using cooperative two photosystems (PSI and PSII). To maintain optimal photosynthetic rates under variable light conditions in nature, phosphorylation of light-harvesting complex for PSII (LHCII) balances the excitation energy distribution between the two photosystems. Here, we investigated the mechanism of this balancing in a green alga. We show that phospho-LHCIIs physically bind to both photosystems. The energy transfer from the LHCIIs to the PSII core complexes becomes less efficient, whereas the excitation level of PSI increases. The time-resolved fluorescence spectra showed an increase in delayed PSI fluorescence, which represents energetical spillover from PSII to PSI. In addition, the spillover is likely mediated by phospho-LHCIIs and PSI antennas. We hypothesize that the spillover explains the larger extent of phospho-LHCIIs dependent energy balancing in the green alga than land plants, which is important for the short-term photoadaptaion in the algal habitat.

## Introduction

In photosynthesis, photosystem I (PSI) and photosystem II (PSII) in the chloroplast thylakoid membrane function in tandem and use light energy to drive redox processes. Under changing light conditions, the excitation levels of these photosystems are balanced to maintain an optimal photosynthetic rate. To this end, plants and green algae have an acclimation mechanism, called the state transition, which redistributes light-harvesting complex II (LHCII) between the two photosystems **(1, 2)**. When PSII is preferentially excited (state 2-inducing conditions), the plastoquinone (PQ) pool gets reduced, which causes plastoquinol to bind to the cytochrome *b*_6_*f* complex (Cyt *b*_6_*f*). Cyt *b*_6_*f* in turn activates a protein kinase (or kinases) required for LHCII phosphorylation **(3)**. The phosphorylated LHCIIs (phospho-LHCIIs) migrate toward PSI to increase the absorption cross-section of PSI under state 2 conditions and thus balance photosystem excitation **(4)**.

Several thylakoid-bound kinases have been reported, including the serine-threonine kinases STT7 and STL1 in *Chlamydomonas reinhardtii* (corresponding to STN7 and STN8 in *Arabidopsis thaliana*, respectively) and <u>t</u>hylakoid <u>a</u>ssociated <u>k</u>inase <u>1</u> (TAK1) in *Arabidopsis* **(5)**. STT7 phosphorylates LHCIIs during state 1 to state 2 transitions and a mutant lacking STT7 was unable to phosphorylate LHCII under state 2-inducing conditions **(6)**. Moreover, several reports indicated that STT7 interacts with subunit IV of Cyt *b*_6_*f* in the stroma region **(7)**. STT7 was activated (via physical modification **(8)**) when plastoquinol bound to the Qo pocket of the Cyt *b*_6_*f* complex **(9)**. Lemeille *et al*. revealed that STT7 activation requires a disulfide bridge between two cysteines at the N-terminus **(10)**. The activated STT7 kinase is thought to directly phosphorylate LHCIIs.

Although many reports have described the roles of LHCII kinases in land plants and green algae, the molecular functions of phospho-LHCIIs during state 2 transitions are still under debate. Kyle *et al*. observed a significant amount of phospho-LHCIIs in the stroma lamellae of the thylakoid membranes, thus increasing the energy transfer between PSI and phospho-LHCIIs **(11)**. Leister and coworkers suggested that phospho-LHCIIs were constantly associated with PSI in an Arabidopsis mutant lacking PsaE, a subunit of PSI **(12)**. These reports agree with the model that phospho-LHCIIs migrate from appressed domains (grana, where PSII is enriched) to non-appressed domains (stroma lamellae, where PSI is enriched) of thylakoids during the state 2 transition, and thus, the phospho-LHCIIs connect to PSI **(13)**. In contrast, Snyders and Kohorn observed significant amounts of LHCIIs in a roughly purified PSI-enriched fraction from a *tak1* mutant, indicating that LHCIIs could associate with PSI even when their phosphorylation was impaired, leading the authors to propose that LHCII phosphorylation was not essential for the lateral migration of LHCII toward PSI **(5)**. Phospho-LHCIIs could associate with PSII in the absence of either the PSI-H or -L subunit in Arabidopsis **(14)**. Tikkanen and colleagues also reported that LHCII phosphorylation was not a prerequisite for physical dissociation of the proteins from the PSII core, but instead contributed to the lateral migration of PSI-LHCII supercomplexes toward the grana margin **(15, 16)**. While CP26, CP29 and LhcbM5 have been reported to bind to PSI during state 2 transitions in C. reinhardtii (Kargul et al. 2005, Takahashi et al. 2006, Drop et al. 2014), CP26 and CP29 are not found in the PSI-LHCI-LHCII supercomplex structure observed by Cryo-EM (Pan et al. 2021). Thus, it remains unclear whether phosphorylated LHCIIs shuttle from PSII to PSI.

Because the green alga *C. reinhardtii* has a large capacity for state transitions **(17)**, the molecular functions of phospho-LHCIIs have been extensively investigated in this species. Iwai *et al*. observed that LHCIIs in PSII-LHCII supercomplexes were not phosphorylated, but those that dissociated from PSII during the induction of state 2 were heavily phosphorylated, leading the authors to hypothesize that phosphorylation is necessary for LHCIIs to undock from PSII **(18)**. The undocked phospho-LHCIIs were shown to dissipate excitation energy, not only increasing the absorption cross-section of PSI during state 1 to state 2 transitions **(19, 20)**. In contrast, Wollman and colleagues reconfirmed that all phospho-LHCIIs that were detached from PSII functionally bound to PSI without dissipating excitation energy, based on the complementary changes in antenna sizes between PSII and PSI during state 1 to state 2 transitions observed by spectroscopic measurements **(17)**. Given the discrepancy over the consequence of state transitions in *C. reinhardtii*, we performed comprehensive *in vivo* and *in vitro* analyses to yield unambiguous insights into the functions of phospho-LHCIIs.

To elucidate the molecular functions of phospho-LHCIIs under state 2 conditions, biochemical and spectroscopic analyses were conducted for cells and photosynthetic supercomplexes stabilized by amphipol **(21)**. Biochemical approaches showed that the STT7-dependent phosphorylation of LHCIIs was not a prerequisite for dissociation from PSII core complexes. By time-resolved fluorescence spectroscopy, we also demonstrated that phospho-LHCIIs altered the energy-transfer dynamics within the PSII-LHCII supercomplexes. Additionally, the time-resolved fluorescence spectra showed an increase in delayed fluorescence from PSI in state 2-induced cells, suggesting a novel function of phospho-LHCIIs whereby energy transfer from PSII-LHCII to PSI-LHCI is stimulated via “spillover”.

## Results

*C. reinhardtii stt7* mutants such as *stt7-1* and *-2* **(6)** or *stt7-9* **(22)** are deficient in the general kinase required for phosphorylation of LHCIIs and therefore can be used to investigate the molecular functions of phospho-LHCIIs. In this study, we generated a single point mutation, named *stt7-o*, via chemical-based random mutagenesis. The mutation in *stt7-o* causes a single amino acid change from glycine to aspartic acid at amino acid 149 (supplemental Figure S1A), which is internal of the kinase consensus domain of STT7. Although the *stt7-o* mutant accumulated almost comparable amount of STT7 protein to WT (supplemental Figure S1B), this mutant showed a severely deficient LHCII-phosphorylation phenotype under state 2 inducible conditions (supplemental Figure S1C). These results indicate that the *stt7-o* mutant harbors STT7 protein without its kinase function. We thus decided to use the *stt7-o* mutant as a mutant deficient in LHCII phosphorylation.

To evaluate the state transition capability of the *stt7-o* mutant, we first observed fluorescence-quenching kinetics under dark-anaerobic conditions where the state 1 to state 2 transition was induced **(17)**. As reported previously for the *stt7-1, -2*, and *-9* mutants, the *stt7-o* mutant was deficient in state transition-dependent fluorescence quenching (qT), whereas the wild-type (WT) strain showed a significant decrease in the maximum fluorescence yield under dark-anaerobic conditions (Figure 1A, also see Figure S2 for fluorescence traces).

**Figure 1.**
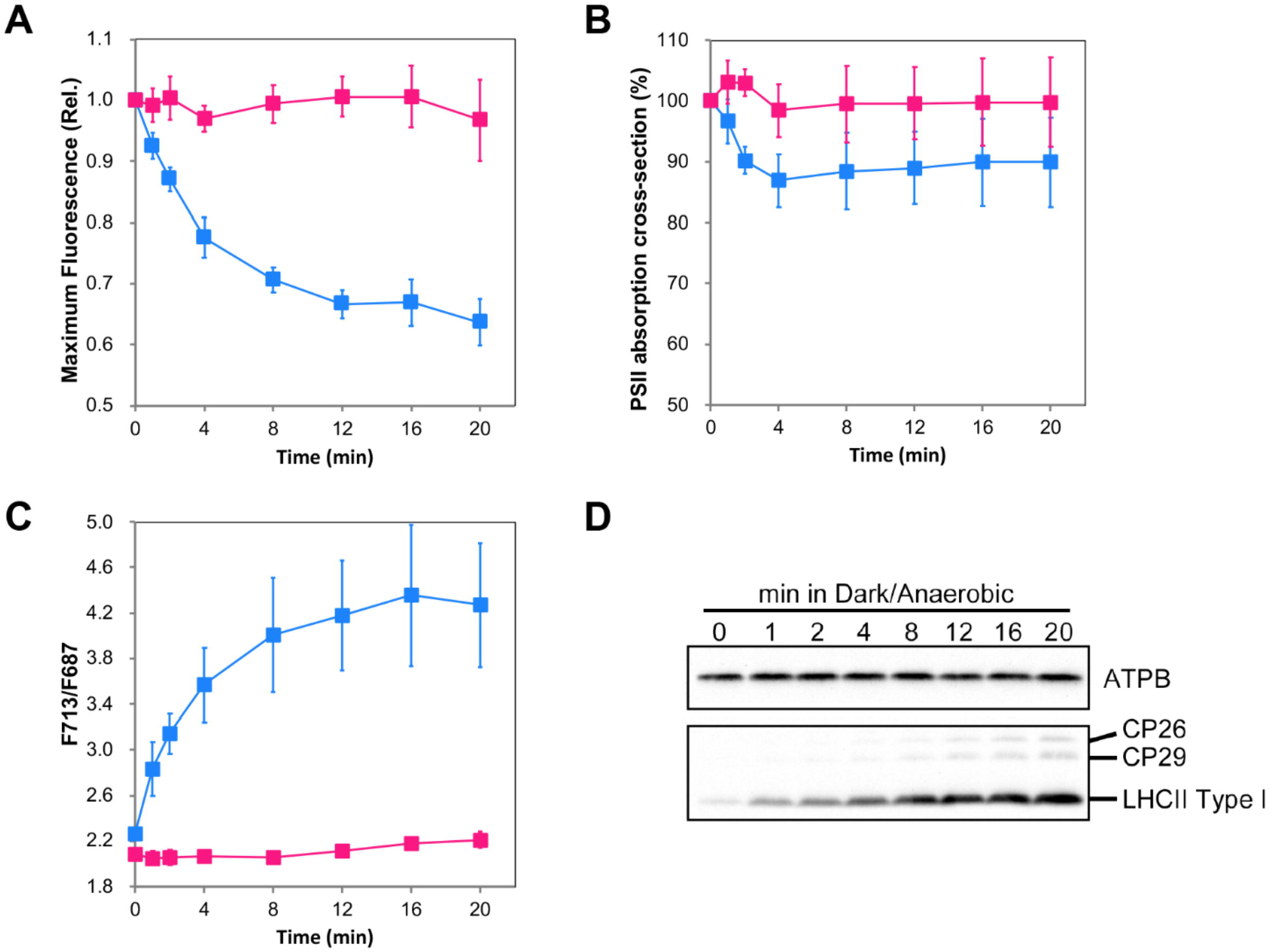
STT7-dependent fluorescence modification under dark-anaerobic conditions. The WT (blue) and *stt7-o* (red) strains were treated under dark-anaerobic conditions for 20 min. (A) The maximum fluorescence yield, (B) relative PSII antenna absorption cross section, and relative PSI (F713) to PSII (F687) fluorescence signal ratio at low-temperature (77 K) were recorded at the same time points. The ratio was normalized to the initial state (state 1 conditions) of the samples. (D) Phosphorylation of LHCII proteins during the measurements is also shown as a representative example of three biological measurements. n = 3 biological replicates; mean ± s.e.

To estimate the amplitude of antenna dissociation from PSII during the state 2 transition, we obtained single-turnover flash-saturation profiles of chlorophyll fluorescence to calculate the relative functional absorption cross-sections of PSII. The measurements of the PSII antenna absorption cross-section during state 1 to state 2 transitions showed a decrease in the maximum fluorescence yield (∼40%; Figure 1A), which is comparable to the level previously reported **(17)**. Interestingly, it is inconsistent with the change of the relative PSII functional antenna size (∼10%, Figure 1B), implying that the LHCIIs were not functionally dissociated from the PSII core, but quenched the excitation energy of PSII.

State transitions involve mechanisms that enable the two photosystems to balance their excitation levels. To test this, we next performed fluorescence spectra analysis under low temperature (77 K) to examine excitation energy re-distribution between the photosystems. Upon functional association of LHCIIs with PSI under state 2 conditions, an increase in the fluorescence peak could be observed at 713 nm because the fluorescence in the ranges of 670– 700 nm and 700–750 nm are attributable to PSII and PSI, respectively. The relative excitation levels, shown as F713/F687 (i.e., the PSI/PSII emission ratio), increased during state 2 transitions (Figures 1C). These data, together with the phosphorylation kinetics of LHCIIs during state 1 to state 2 transitions (Figure 1D), suggest that the LHCIIs were not functionally dissociated from PSII, although they were phosphorylated and re-distributed energy from PSII to PSI.

An inconsistency between the functional PSII antenna size and the fluorescence quenching in state 2 implied that most phospho-LHCIIs were not physically detached from PSII, even in WT cells. To further evaluate the fate of phospho-LHCIIs, we next performed sucrose density gradient purification of the photosystems from state 2-locked thylakoid membranes. In these membranes, LHCIIs were heavily phosphorylated (Figure S3), and showed quenching capability as demonstrated by faster fluorescence decay compared to *stt7-o* thylakoids (Figure S4). We obtained four different fractions from WT membranes, corresponding to free LHCII, PSI-LHCI, PSI-LHCI/II, and PSII-LHCII supercomplexes from top to bottom of the tube (Figure 2A). No PSI-LHCI/II band was observed in *stt7-o* thylakoids. As expected, the PSII-LHCII supercomplex fraction appeared to be equally distributed in the state 2-induced membranes from the WT and *stt7-o* strains.

**Figure 2.**
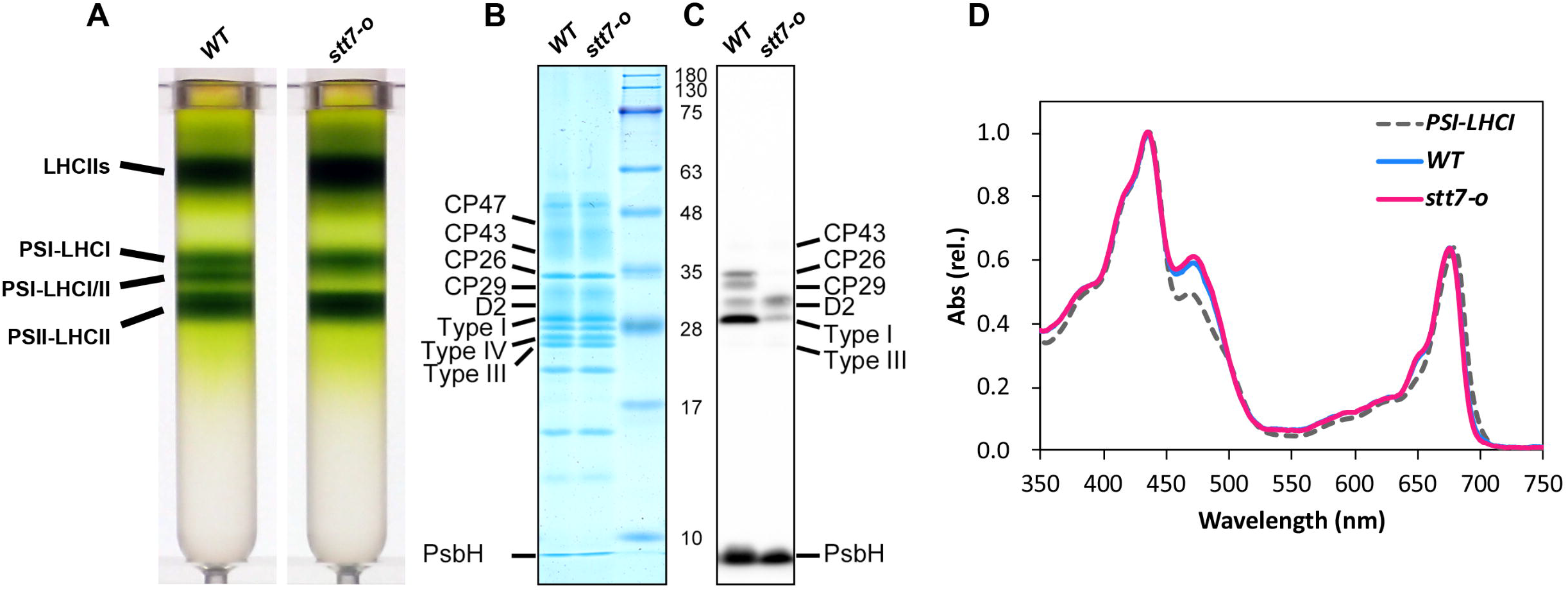
Purification and characterization of PSII-LHCII supercomplexes under state 2 conditions. (A) Sucrose gradient fractionation of solubilized state 2 thylakoid membranes. Each band shown is labeled. (B) SDS-PAGE analysis of purified PSII-LHCII supercomplexes from WT and *stt7-o* cells. Sample loading was normalized to 2.5 μg Chl/lane. (C) Immunoblotting analysis performed to visualize phosphorylated polypeptides in the supercomplexes. Phosphorylated proteins were labeled according to SDS-PAGE analysis, as described in a previous study **(47)**. (D) Absorption spectra of the supercomplexes obtained from WT (blue) and *stt7-o* (red) strains normalized at the peak of the Soret region. The spectrum of PSI-LHCI is shown as a control.

The PSII-LHCII supercomplex fraction in the WT strain contained the same LHCII composition as in the *stt7-o* mutant, as revealed by sodium dodecyl sulfate-polyacrylamide gel electrophoresis (SDS-PAGE) analysis (Figure 2B), but the phosphorylation levels of LHCIIs were clearly different (Figure 2C). Indeed, densitometric analysis of phospho-immunoblot images revealed that at least four-fold heavier phosphorylation occurred in WT LHCIIs than in *stt7-o* LHCIIs including CP26, CP29, and LHCII type I (Figure S5). The absorption spectra of those PSII-LHCII fractions also showed superimposable shapes, even between 470 and 650 nm (Figure 2D), which represents chlorophyll *b* absorption that is mainly coordinated in LHCIIs **(23, 24)**. These data suggested that the amount of LHCIIs associated with PSII was almost identical in the PSII-LHCII supercomplexes obtained from the WT and *stt7-o* strains.

Steady-state fluorescence spectra of the PSII-LHCII supercomplexes at low-temperature (77 K) showed negligible fluorescence from free LHCIIs (shoulder around 680 nm), even with the preferential excitation of chlorophyll *b* (LHCII) at 480 nm (Figure S6). These results further confirmed that the phospho-LHCIIs remained physically and tightly associated with the PSII core complexes to absorb light energy, similarly in both strains under state 2 conditions. The increase of fluorescence intensities in the ranges of 700–720 nm, which were only detected in PSII complexed with phospho-LHCIIs (Figure S6; blue line, WT) implies the conformational changes of LHCIIs as reported in higher plants **(25, 26)**. Because the PSII-LHCII supercomplexes were not contaminated with PSI complexes as shown by SDS-PAGE analysis (Fig. 2B), it is plausible that the increase of 700-720 nm fluorescence intensities implies modifications in the conformation of the PSII-LHCII supercomplexes due to LHCII phosphorylation.

Fluorescence quenching analysis (Figure 1) showed that phosphorylated LHCIIs in PSII (Figure 2) contributed to fluorescence quenching under state 2 conditions *in vivo*. To investigate energy-transfer dynamics in PSII-LHCII supercomplexes with (WT) or without (*stt7-o*) phospho-LHCIIs, we performed time-resolved fluorescence measurements followed by fluorescence decay-associated spectra (FDAS) analysis (gray lines in Figures 3 and S7, Table S1). In the 1st component of FDAS (∼65 ps), we observed the energy transferred from LHCIIs to the PSII core antenna CP43 as indicated by major positive peak around 680 nm (LHCII) and negative peak around 685 nm (CP43). The red 700–720 nm component in the PSII-LHCII supercomplex of WT was also observed, and was pronounced at the energy-trapping (∼150 ps) and equilibration states of the PSII core (∼600 ps), indicating that the phospho-LHCII-dependent formation of a low-energy state was coupled with PSII core complexes.

**Figure 3.**
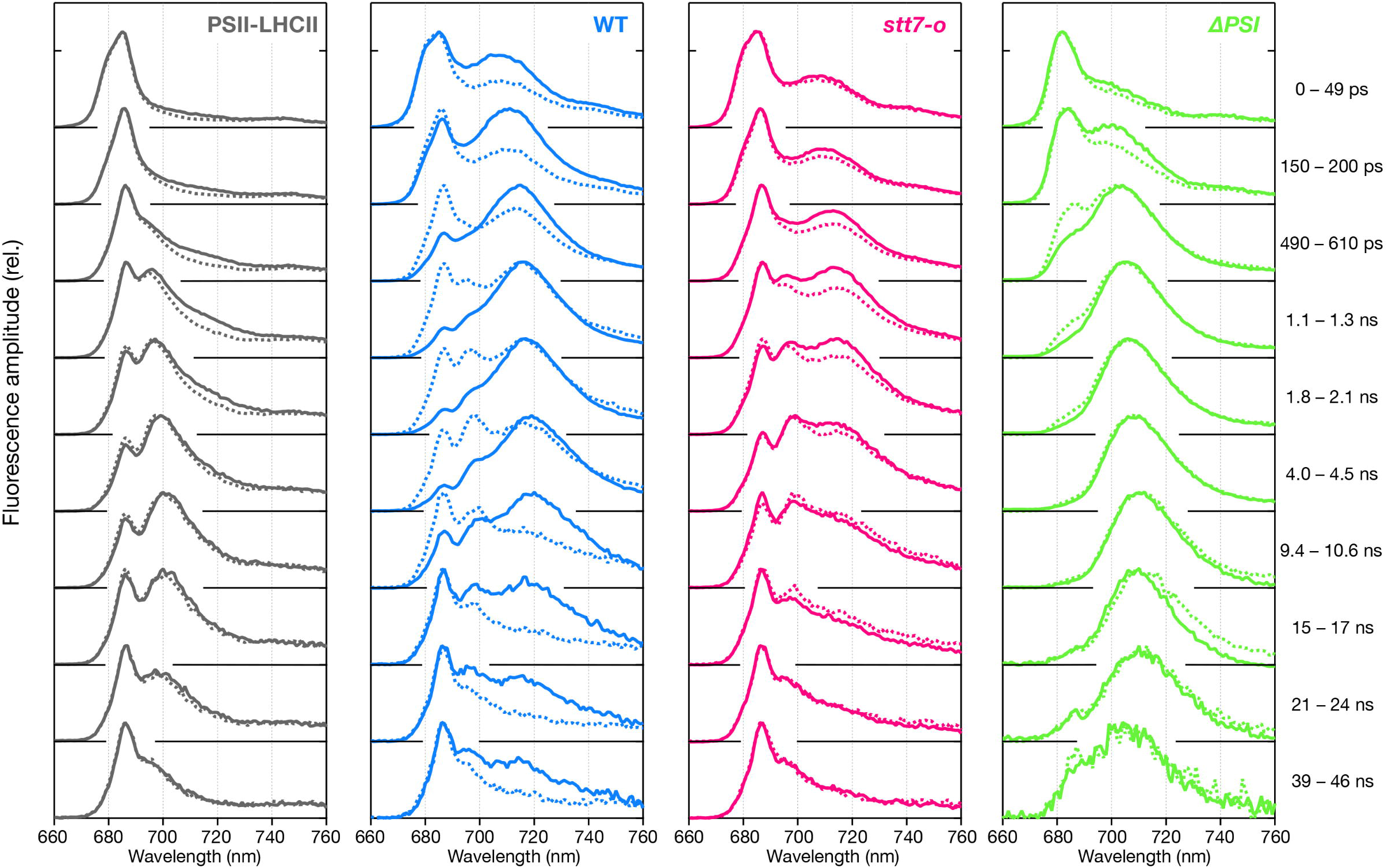
Time-resolved fluorescence spectra of the isolated PSII-LHCII supercomplexes and *Chlamydomonas* cells at 77 K. Time-resolved fluorescence spectra of the PSII-LHCII supercomplexes (gray) were obtained under state 2 conditions from WT (solid line) and *stt7-o* strains (dashed line) at 77 K. WT (blue) and *stt7-o* (red) cells under state 1 (dashed line) and state 2 (solid line) conditions were also measured by 459 nm excitation at 77 K.

To qualitatively evaluate whether the altered excitation energy dynamics contributed to excitation energy quenching, we performed up-conversion time-resolved fluorescence analysis of the PSII-LHCII supercomplexes at room temperature. This analysis revealed only a little quenching in the isolated PSII-LHCII supercomplexes (Figure S8), regardless of whether the LHCIIs were phosphorylated. This result, together with the obvious fluorescence quenching in both cell types (Figure 1A) and isolated thylakoids (Figure S4) in state 2, could indicate the presence of a fluorescence quencher in the membrane that was not tightly associated with PSII supercomplexes.

Based on these observations, the question arises as to why PSII-LHCII supercomplexes alter the energy-transfer dynamics following LHCII phosphorylation. In other words, it is unclear how phospho-LHCIIs in PSII-LHCII supercomplexes contribute to the excitation energy redistribution between photosystems in state 2. The results obtained above implied that the phospho-LHCIIs in the supercomplexes were involved in direct energy transfer from PSII-LHCII to PSI-LHCI, which is traditionally termed “spillover”. To verify the energy transfer occurring between the membrane-embedded photosystems, we obtained 77 K time-resolved fluorescence spectra of WT and *stt7-o* cells under both state 1 and state 2 conditions (Figure 3). After the transition from state 1 to state 2, the PSI fluorescence (710–720 nm) in WT cells was enhanced from the initial phase (0–49 ps) to 17 ns after excitation, representing an efficient redirection of the energy to PSI, rather than PSII, whereas *stt7-o* mutant showed negligible change in this fluorescence (Figure 3). This phenomenon corresponds to the re-association of LHCIIs with PSI and the formation of PSI-LHCI-LHCII supercomplexes (Figure 2A), representing a conventional state 2 transition.

We next analyzed the delayed fluorescence, which shows longer lifetime than intrinsic relaxation of chlorophylls. Because delayed florescence with a time constant of ∼20 ns at 77 K is specifically derived from the charge recombination in the PSII reaction center **(27)**, the delayed fluorescence spectrum reflects energy migration from the PSII core complexes. After the transition from state 1 to state 2, the delayed fluorescence spectra in WT cells showed relatively increased fluorescence in the PSI-fluorescence region (around 717 nm), whereas *stt7-o* cells showed no obvious increase (red traces in Figure 4 and Table S2, WT and *stt7-o*). The increased amplitude of the delayed fluorescence in the PSI-fluorescence region indicates that the energy migrated from PSII core complexes to PSI under state 2 conditions.

**Figure 4.**
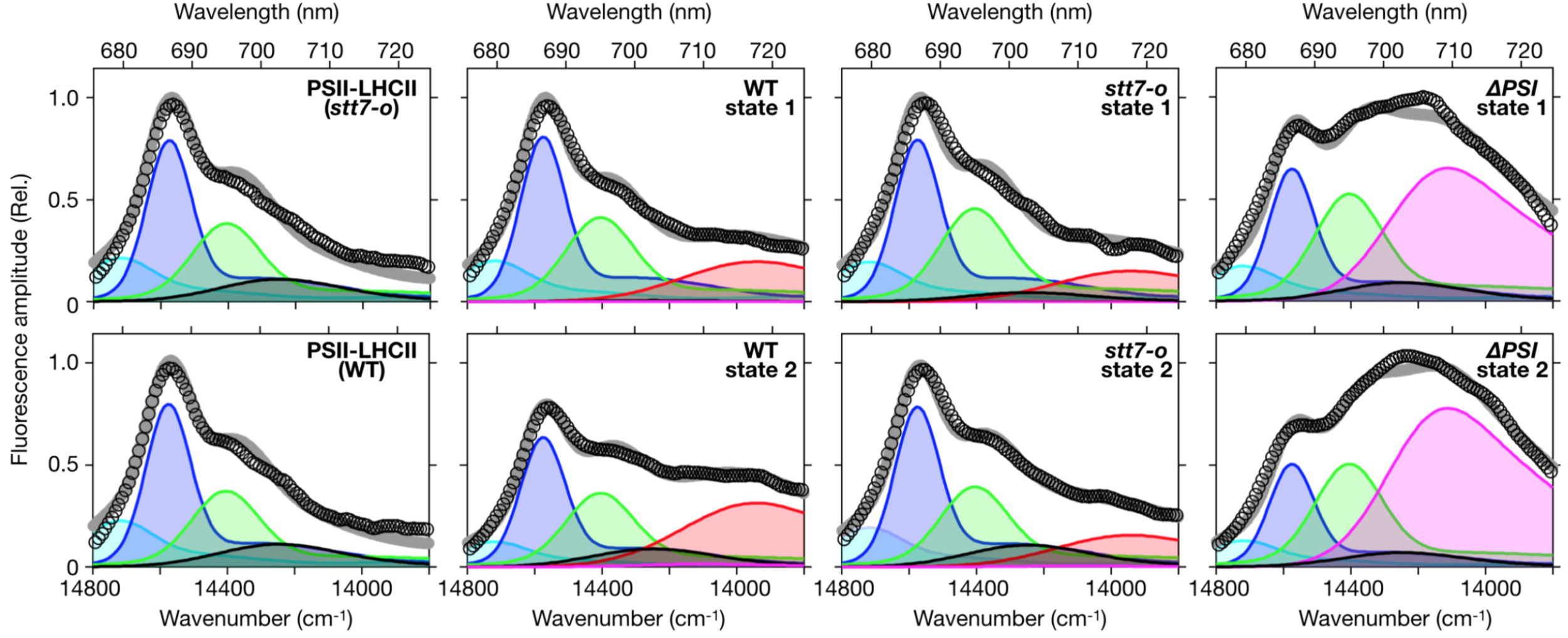
Delayed fluorescence spectra of the isolated PSII-LHCII supercomplexes and *Chlamydomonas* cells obtained at 77 K. FDAS of delayed fluorescence obtained from state 2 PSII-LHCII supercomplexes (Upper; WT, Lower; *stt7-o*,) and cells under state 1 (upper panel) and state 2 (lower panel) conditions were decomposed into six components (details shown in Table 1). Open circles and gray lines represent FDAS spectra and fitting curve, respectively. The peaks and width of components were obtained by global fitting based on wavenumber (cm^−1^). Each color of component peaks represents 679 nm (cyan, LHCII), 686 nm (blue, CP43), 694 nm (green, CP47), 702 nm (black, phospho-LHCII and/or phospho-PSII), 709 nm (magenta, LHCI) and 717 nm (red, PSI-LHCI). The fluorescence amplitudes were normalized with the total delayed fluorescence intensity of each samples. Complete FDAS results are shown in Figure S7.

This interesting change in the delayed fluorescence suggests that the transition from state 1 to state 2 not only induced the migration of LHCII from PSII to PSI, but also facilitated energy transfer from the PSII core to PSI. Indeed, the estimated ratio of spillover from the PSII-PSI complex to total PSII, which was calculated as in a previous report **(28)**, increased two-fold from 9.8% in state 1 to 19% in state 2 in WT cells. In addition, the results of a decomposition fitting into the delayed fluorescence spectra clearly showed that the amplitudes of LHCII (679 nm) and PSII (CP43; 686 nm, CP47; 694 nm) decreased, whereas the amplitude of PSI (717 nm) and possibly phospho-LHCII and/or phospho-PSII (702 nm) increased in state 2-induced WT cells (Figure 4 and Table S2). These results imply that phospho-LHCIIs under state 2 conditions induce energy transfer from PSII to PSI.

To investigate the energy interaction between PSII and PSI, we also analyzed the delayed fluorescence in a mutant deficient in the PSI core (Δ*PSI*) (Figure S9). Although no decrease in maximum fluorescence yield was observed during the state 1 to state 2 transition (Figure S9A), state 2-inducible LHCII phosphorylation (Figure S9B) and changes in delayed fluorescence spectra were observed in the Δ*PSI* strain (Figures 4 and Table S2). The delayed fluorescence spectra showed a relatively large peak at 709 nm, which probably originated from LHCI (magenta traces in Figure 4 and Table S2). These findings may represent an energy transfer from the phosphorylated PSII-LHCII to free LHCI in the Δ*PSI* strain, implying that energy transfer between photosystems was mediated by LHCI. Time-resolved fluorescence analysis of purified PSII-LHCII supercomplexes obtained from WT and *stt7-o* strains under state 2 conditions showed almost identical spectra in delayed fluorescence (Figure 4), indicating that energy was maintained in PSII-LHCII when PSI core and/or LHCI was absent. This further supported the idea that the delayed florescence peak at 709 nm observed in Δ*PSI* mutant cells were generated via spillover-dependent energy transfer from PSII-LHCII to LHCI. Collectively, these results indicate that STT7-dependent LHCII phosphorylation regulates direct energy interactions or “spillover” between PSII-LHCII and PSI-LHCI supercomplexes.

## Discussion

State transitions have been recognized as energy balancing mechanisms mediated by LHCII shuttling between photosystems. The phosphorylation of LHCIIs is a prerequisite for this energy redistribution in thylakoid membranes. However, no reports have unambiguously demonstrated the lateral migration of phosphorylated LHCIIs during state transitions. Here, we present results showing that phosphorylated LHCIIs were not substantially dissociated from PSII in a phosphorylation-dependent manner (Figure 2). However, the increased energy accumulation in PSI was shown as an increased fluorescence ratio (F713/F687) in WT cells (Figure 1). This result confirms the presence of phospho-LHCII-dependent excitation energy redistribution between photosystems, which is widely accepted **(29)**. Time-resolved fluorescence spectra analysis revealed that the redistribution of energy was coupled with energy transfer from the PSII core to PSI-LHCI supercomplexes. These observations indicated that the molecular mechanisms underlying the phospho-LHCII-dependent increase in PSI excitation involve spillover from PSII to PSI.

It is difficult to distinguish between the phenomena of state transitions and spillover because both mechanisms contribute similarly to energy redistribution between photosystems. Indeed, many researchers have concluded that phospho-LHCII-dependent energy redistribution reflected either state transitions or spillover (reviewed in **(13, 30)**). A consensus of the mechanistic significance of STT7-dependent LHCII phosphorylation has not been reached thus far. In this study, we focused on delayed fluorescence spectra (Figures 4 and S7) to evaluate whether the STT7-dependent energy redistribution was mediated by the migration of phospho-LHCIIs from PSII to PSI and/or via another mechanism involving phospho-LHCIIs. Delayed fluorescence represents re-excitation of chlorophylls after charge recombination in the PSII reaction center, where the fluorescence is not derived from other components, such as free LHCII or PSI **(28)**. The delayed fluorescence observed in WT cells showed state 2-inducible spillover from PSII-LHCII to PSI-LHCI, which was represented by strengthened delayed fluorescence from 700–720 nm (Figure 4 and Table S2, WT). To our knowledge, this study represents the first observations that clarify the contribution of STT7-dependent LHCII phosphorylation to spillover in green algae.

The question arises as to how phospho-LHCIIs contribute to energy migration from PSII to the PSI reaction center. PSII-LHCII supercomplexes, which have phospho-LHCIIs, showed obvious red fluorescence from 700–710 nm (Figures 3 and S6) that were probably due to conformational changes induced by phosphorylation **(31)**. The occurrence of red fluorescence may allow migration of energy from the PSII core to phospho-LHCIIs. Intriguingly, however, delayed fluorescence was also observed in the 709 nm peak region, which represents the fluorescence from LHCIs in the mutant with a deficiency in the PSI core (Figures 4 and S7), suggesting that LHCIs accept energy from PSII-LHCII. Based on these observations, it is possible that an energy-transfer pathway exists between phospho-LHCIIs and LHCIs.

Phospho-LHCIIs have been shown to localize around the PsaH subunit in state 2 in a green alga **(32, 33)**, where green algal specific LHCIs (LHCA2 and LHCA9) are predicted to be located **(34, 35)**. The possible interaction between phospho-LHCIIs and LHCA2 and/or LHCA9 may contribute to spillover under state 2 conditions. On the other hand, because several LHCIs showed a fluorescence peak near to 709 nm region **(36)**, it was also plausible that those LHCIs (possibly LHCA4, LHCA5, LHCA6, and LHCA8) might contribute to spillover mechanism. Further comprehensive study, including genetic, biochemical, structural, and spectroscopic analyses of the *lhca* mutants, is required to obtain a detailed molecular characterization of the spillover mechanism.

Spillover involves a mechanism that uses PSI as a fluorescence quencher to avoid PSII over-excitation **(37)**. Recently, we also discovered light-<u>h</u>arvesting <u>c</u>omplex <u>s</u>tress related 1 (LHCSR1)-mediated fluorescence quenching in PSI **(38)**. Algae expressing LHCSR1 under ultraviolet radiation showed large fluorescence quenching via increased energy transfer from LHCIIs to PSI. The difference between the spillover and LHCSR1-mediated fluorescence quenching in PSI is defined by pH dependency. The former mechanism does not require a low pH to activate fluorescence quenching, whereas the latter mechanism does. Further, spillover is independent of *de novo* synthesis of quencher proteins such as PsbS, LHCSR1, and LHCSR3. For these reasons, phospho-LHCII-dependent spillover likely serves as a primary acclimatizing response to various environmental conditions. STT7-dependent LHCII phosphorylation is regulated through plastoquinone reduction in response to both light and anoxia **(3)**. The anoxic condition inhibits respiration, which in turn reduces the plastoquinone pool by increasing the total reducing power within cells. Because *Chlamydomonas* species live in soils dominated by bacterial inhabitants **(39)**, algal cells are often exposed to anoxic conditions, particularly during dusk, when photosynthesis is slow. To avoid PSII over-excitation following sunrise, green algae should be able to utilize the phospho-LHCII-dependent spillover system. It is reasonable to expect that the energy redistribution to PSI via spillover could protect PSII, considering that the system is a rapid (assembled without light) and low-cost (no *de novo* protein synthesis required) photoacclimation mechanism. Thus, the spillover system in green algae is likely optimized for survival in native habitats.

It appears that spillover is one of several widely conserved photoacclimation mechanisms operating in photosynthetic organisms. Aro and colleagues showed that spillover occurred at the grana margin of the thylakoid membrane in land plants **(15, 16)**. Yokono *et al*. also found the spillover megacomplex in thylakoid membranes from Arabidopsis **(28)**. Here, we discovered phospho-LHCII-mediated spillover in a green alga, whose thylakoid membranes are loosely appressed compared to the grana of land plants **(40)**. The spillover mechanism was also found in red lineages, including a red alga **(28)** and *Symbiodinium* **(41)**. Spillover also occurs in cyanobacteria, which lack chloroplasts and the grana structure of thylakoids **(42-44)**, implying that this mechanism was established at a very primitive stage in the evolution of photosynthesis. Thylakoid membrane structures **(44)** and light-harvesting antennas **(45)** are widely diversified among these photosynthetic organisms: however, it seem that photosynthetic organisms have retained this simple and effective photoprotection mechanism, “spillover”, which mainly involves photosystem core complexes, over the course of their evolution.

## Conclusion

The findings of the present study link the conventional molecular mechanism of STT7-dependent state transitions and spillover. In *Chlamydomonas*, phosphorylated LHCIIs mediate facile transfer of energy from PSII-LHCII to PSI-LHCI, which is recognized as spillover. The presence of phospho-LHCII-dependent spillover was first hypothesized over 30 years ago **(30)**; however, this hypothesis was refuted by studies emphasizing phospho-LHCII migration between the photosystems **(13)**. The model presented in this study (Figure 5) overcomes the contradiction between the previous reports because the phospho-LHCIIs mediate both the attachment of LHCII to PSI (state transitions) and direct energy transfer from PSII-LHCII to PSI-LHCI (spillover).

**Figure 5.**
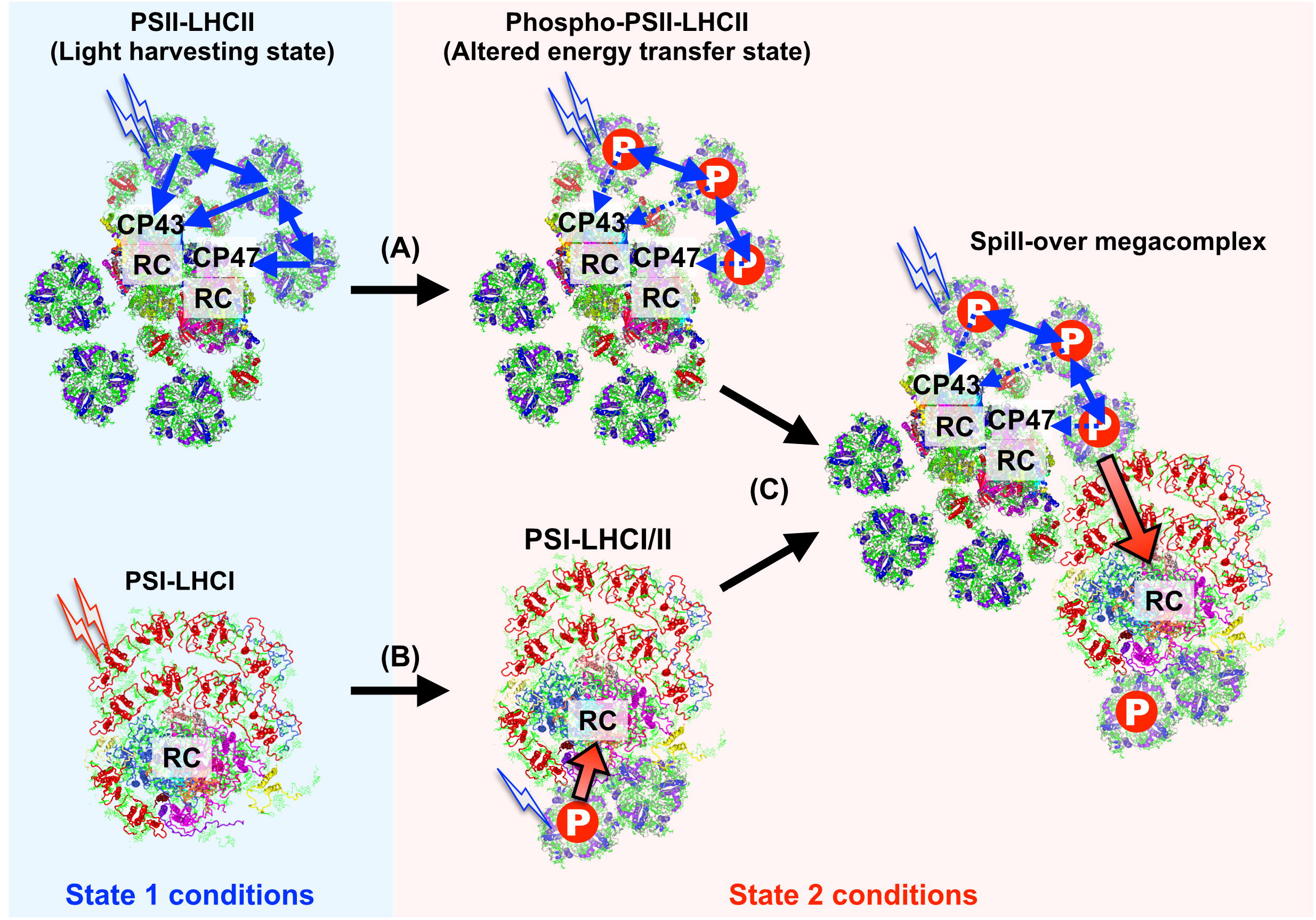
Phospho-LHCII regulated spillover between photosystems. (A) STT7-dependent phosphorylation under state 2 conditions alters the excitation energy transfer between LHCIIs and PSII. (B) Meanwhile, the PSI-LHCI supercomplex associates with LHCIIs (conventional state 2 transition). (C) The PSI-LHCI(-LHCII) supercomplex approaches the phosphorylated PSII-LHCII supercomplex to form a spillover megacomplex to efficiently transfer the excitation energy accumulated in PSII to PSI probably via LHCI.

## Materials and Methods

### Culture conditions

The *C. reinhardtii* strain 137c (mt+) was obtained from the Chlamydomonas Center (http://www.chlamy.org) and was used as the WT strain. The mutant strain *stt7-o* was isolated after performing random mutagenesis. For the random mutagenesis, 1-methyl-3-nitro-1-nitrosoguanidine (Wako, Japan) was used at a final concentration of 5 μg/mL, according to previous reports **(39, 46)**. We screened the mutants, for those were unable to perform qT quenching (state 2 transitions), using a FluoCAM 800MF (Photon System Instruments, Czech Republic). The mutated locus in the *stt7-o* strain was identified by genome sequencing after genetic mapping (Figure S1A), and the obtained *stt7-o* mutant was then backcrossed four times with the WT strain. After the backcross process, we roughly characterize the phenotype through immunoblotting and 77 K fluorescence spectra analysis (Figure S1B-D). The Δ*PSI* (Δ*PsaA*) strain was obtained as described previously **(47)**. All strains were grown in Tris-acetate-phosphate (TAP) medium **(39)** under dim light (< 20 μE m^−2^ s^−1^) at 23°C until they reached the mid-log growth phase. The culture medium was exchanged with high-salt (HS) minimal medium with slight modifications **(39, 48)** 16 hours before the experiments. In all experiments, the cultured cells were induced to become state 1 with dim far-red light (< 5 μE m^−2^ s^−1^) at 23°C for 30 min prior to the state 2 induction treatment (dark-anaerobic conditions as previously described **(17)**).

### Isolation of thylakoid membranes and photosynthetic supercomplexes

Thylakoid membranes from the WT and *stt7-o* strains after state 1 or state 2 inductions were isolated as previously reported **(49)**. To purify the photosynthetic supercomplexes from the isolated thylakoids, we used amphipol A8-35, as follows. Thylakoid membranes were detergent-solubilized as described in a previous report **(47)**, then immediately mixed with A8-35 at a final concentration 1.0%, and incubated on ice for 15 minutes. The A8-35-treated membranes were separated by detergent-free sucrose density gradient ultracentrifugation at 230,000 x g for 16 hours.

### SDS-PAGE and immunoblotting analysis

SDS-PAGE and immunoblotting analyses were performed as described previously **(50)**. For the phosphorylation analysis, Phospho-Threonine antibody was obtained from Invitrogen (71-8200, rabbit polyclonal), and used as in a previous study **(49)**. Antisera against ATPB was obtained from Agrisera (AS05 085, rabbit polyclonal); antisera against STT7 was raised against synthetic peptide corresponding to 695-713 amino acid of STT7 mature protein. For relative quantification of the phosphorylated proteins, images were analyzed using Image Lab 5.0 software (Bio-Rad, Hercules, CA), and the obtained band intensities were normalized to the intensity of the control samples indicated in the figure legends.

### Spectroscopic fluorescence measurements

Fluorescence quenching kinetics were measured with a PAM-2500 instrument (Walz, Germany) under dark-anaerobic conditions as previously described **(17)**. The functional absorption cross-section that reflects the functional antennae sizes of PSII was measured using a FIRe fluorometer system (Model FIRe; Satlantic, Nova Scotia, Canada), as described previously **(49)**. Low-temperature (77 K) fluorescence spectra were obtained with a FluoroMax4 (HORIBA Jobin-Yvon) as described in a previous study **(51)**. Time-resolved fluorescence decays at room temperature (23°C) were obtained via time-correlated single-photon counting of fluorescence using a FluoroCube (HORIBA Jobin-Yvon) with slight modifications from a previous study **(51)**. A picosecond pulse diode laser (DD-470L; HORIBA Jobin-Yvon) was used to excite chlorophylls at 463 nm with a 10 MHz repetition rate (1.64 pJ/pulse), and emission was detected at 682 nm using a monochromatic detector (bandwidth = 8 nm). To conduct FDAS, time-resolved fluorescence decays at 77 K were obtained by time-correlated single-photon counting of fluorescence as described previously **(28)**. Spectral decomposition was basically performed as described previously **(28)**. In brief, each spectral component is represented as a sum of Gaussian functions. Each spectral component has the same position and width in all samples, and only the amplitude of the spectral components can differ between samples.

## Acknowledgments

We thank Mrs. Rie Hoshi for providing technical support throughout the entire project. We also thank Mrs. Harumi Yonezawa and Mrs. Tamaka Kadowaki for providing technical assistance with mutant isolation of the alga. Dr. Kenji Takizawa is thanked for critical reading of the manuscript. This work was supported by the JSPS KAKENHI (JP15H05599, 20H03282, and 21K19282 to R.T., and JP16H06553 to S.A. and J.M.) and the NINS program for cross-disciplinary study (Grant Number 01311701) to R.T.

## Author contributions

RT and JM conceived the research. RT designed the experiments, and performed the biochemical and physiological analyses. RT, EK, and SA performed the spectroscopic analysis. EK, SA, MY, and YU analyzed the spectroscopic data. KF-K, and NO isolated the *stt7-o* mutant. RT wrote the manuscript with input from all authors. JM supervised the entire research, provided resources, and critically revised the manuscript. All authors contributed to interpretation of the results, and approved the final version of the manuscript.

## Competing interests

The authors declare no competing interests.

## Notes

### Competing Interest Statement

The authors have declared no competing interest.

## References

1. C. Bonaventura, J. Myers, Fluorescence and oxygen evolution from Chlorella pyrenoidosa. Biochim. Biophys. Acta. 189, 366–383 (1969).

2. N. Murata, Control of excitation transfer in photosynthesis I. Light-induced change of chlorophyll a fluoresence in Porphyridium cruentum. Biochim. Biophys. Acta. 172, 242–251 (1969).

3. F.-A. Wollman, State transitions reveal the dynamics and flexibility of the photosynthetic apparatus. EMBO J. 20, 3623–3630 (2001).

4. J. Minagawa, State transitions--the molecular remodeling of photosynthetic supercomplexes that controls energy flow in the chloroplast. Biochim. Biophys. Acta. 1807, 897–905 (2011).

5. S. Snyders, B. D. Kohorn, Disruption of thylakoid-associated kinase 1 leads to alteration of light harvesting in Arabidopsis. J. Biol. Chem. 276, 32169–32176 (2001).

6. N. Depège, S. Bellafiore, J. D. Rochaix, Role of chloroplast protein kinase Stt7 in LHCII phosphorylation and state transition in Chlamydomonas. Science 299, 1572–1575 (2003).

7. L. Dumas et al., A stromal region of cytochrome b6f subunit IV is involved in the activation of the Stt7 kinase in Chlamydomonas. Proc. Natl. Acad. Sci. U. S. A. 114, 12063–12068 (2017).

8. S. K. Singh et al., Trans-membrane signaling in photosynthetic state transitions: redox-and structure-dependent interaction in vitro between STT7 kinase and the cytovhrome b6f complex. J. Biol. Chem. 291, 21740–21750 (2016).

9. G. Finazzi, F. Zito, R. P. Barbagallo, F. A. Wollman, Contrasted effects of inhibitors of cytochrome b_6_f complex on state transitions in Chlamydomonas reinhardtii: the role of Qo site occupancy in LHCII kinase activation. J. Biol. Chem. 276, 9770–9774 (2001).

10. S. Lemeille et al., Analysis of the chloroplast protein kinase Stt7 during state transitions. PLoS Biol. 7, e45 (2009).

11. D. J. Kyle, L. A. Staehelin, C. J. Arntzen, Lateral mobility of the light-harvesting complex in chloroplast membranes controls excitation energy distribution in higher plants. Arch. Biochem. Biophys. 222, 527–541 (1983).

12. P. Pesaresi et al., A stable LHCII-PSI aggregate and suppression of photosynthetic state transitions in the psae1-1 mutant of Arabidopsis thaliana. Planta 215, 940–948 (2002).

13. J. F. Allen, J. Forsberg, Molecular recognition in thylakoid structure and function. Trends Plant Sci. 6, 317–326 (2001).

14. S. Zhang, H. V. Scheller, Light-harvesting complex II binds to several small subunits of photosystem I. J. Biol. Chem. 279, 3180–3187 (2004).

15. M. Tikkanen et al., Phosphorylation-dependent regulation of excitation energy distribution between the two photosystems in higher plants. Biochim. Biophys. Acta. 1777, 425–432 (2008).

16. M. Grieco, M. Suorsa, A. Jajoo, M. Tikkanen, E. M. Aro, Light-harvesting II antenna trimers connect energetically the entire photosynthetic machinery - including both photosystems II and I. Biochim. Biophys. Acta. 1847, 607–619 (2015).

17. W. J. Nawrocki, S. Santabarbara, L. Mosebach, F. A. Wollman, F. Rappaport, State transitions redistribute rather than dissipate energy between the two photosystems in Chlamydomonas. Nature Plants 2, 7 (2016).

18. M. Iwai, Y. Takahashi, J. Minagawa, Molecular remodeling of photosystem II during state transitions in Chlamydomonas reinhardtii. Plant Cell 20, 2177–2189 (2008).

19. M. Iwai, M. Yokono, N. Inada, J. Minagawa, Live-cell imaging of photosystem II antenna dissociation during state transitions. Proc. Natl. Acad. Sci. U. S. A. 107, 2337–2342 (2010).

20. C. Ünlü, B. Drop, R. Croce, H. van Amerongen, State transitions in Chlamydomonas reinhardtii strongly modulate the functional size of photosystem II but not of photosystem I. Proc. Natl. Acad. Sci. U. S. A. 111, 3460–3465 (2014).

21. A. Watanabe, E. Kim, R. N. Burton-Smith, R. Tokutsu, J. Minagawa, Amphipol-assisted purification method for the highly active and stable photosystem II supercomplex of Chlamydomonas reinhardtii. FEBS Lett. 593, 1072–1079 (2019).

22. P. Cardol et al., Impaired respiration discloses the physiological significance of state transitions in Chlamydomonas. Proc. Natl. Acad. Sci. U. S. A. 106, 15979–15984 (2009).

23. Z. Liu et al., Crystal structure of spinach major light-harvesting complex at 2.72 A resolution. Nature 428, 287–292 (2004).

24. J. Standfuss, A. C. Terwisscha van Scheltinga, M. Lamborghini, W. Kühlbrandt, Mechanisms of photoprotection and nonphotochemical quenching in pea light-harvesting complex at 2.5 Å resolution. EMBO J. 24, 919–928 (2005).

25. H. Kirchhoff, H. J. Hinz, J. Rosgen, Aggregation and fluorescence quenching of chlorophyll a of the light-harvesting complex II from spinach in vitro. Biochim. Biophys. Acta. 1606, 105–116 (2003).

26. C. Ilioaia, M. P. Johnson, P. Horton, A. V. Ruban, Induction of efficient energy dissipation in the isolated light-harvesting complex of photosystem II in the absence of protein aggregation. J. Biol. Chem. 283, 29505–29512 (2008).

27. M. Mimuro et al., Delayed fluorescence observed in the nanosecond time region at 77 K originates directly from the photosystem II reaction center. Biochim. Biophys. Acta. 1767, 327–334 (2007).

28. M. Yokono, A. Takabayashi, S. Akimoto, A. Tanaka, A megacomplex composed of both photosystem reaction centres in higher plants. Nat. Commun. 6, 6 (2015).

29. J. D. Rochaix, Role of thylakoid protein kinases in photosynthetic acclimation. FEBS Lett. 581, 2768–2775 (2007).

30. J. Barber, Influence of surface-charges on thylakoid structure and function. Annu. Rev. Plant Physiol. Plant Mol. Biol. 33, 261–295 (1982).

31. H. Zer et al., Regulation of thylakoid protein phosphorylation at the substrate level: Reversible light-induced conformational changes expose the phosphorylation site of the light-harvesting complex II. Proc. Natl. Acad. Sci. U. S. A. 96, 8277–8282 (1999).

32. Z. H. Huang et al., Structure of photosystem I-LHCI-LHCII from the green alga Chlamydomonas reinhardtii in State 2. Nat. Commun. 12 (2021).

33. X. W. Pan et al., Structural basis of LhcbM5-mediated state transitions in green algae. Nature Plants 7, 1119-+ (2021).

34. S.-I. Ozawa et al., Configuration of Ten Light-Harvesting Chlorophyll a/b Complex I Subunits in Chlamydomonas reinhardtii Photosystem I. Plant Physiol. 178, 583–595 (2018).

35. H. Kubota-Kawai et al., Ten antenna proteins are associated with the core in the supramolecular organization of the photosystem I supercomplex in Chlamydomonas reinhardtii. J. Biol. Chem. 294, 4304–4314 (2019).

36. M. Mozzo et al., Functional analysis of photosystem I light-harvesting complexes (Lhca) gene products of Chlamydomonas reinhardtii. Biochim. Biophys. Acta. 1797, 212–221 (2010).

37. A. Derks, K. Schaven, D. Bruce, Diverse mechanisms for photoprotection in photosynthesis. Dynamic regulation of photosystem II excitation in response to rapid environmental change. Biochim. Biophys. Acta. 1847, 468–485 (2015).

38. K. Kosuge et al., LHCSR1-dependent fluorescence quenching is mediated by excitation energy transfer from LHCII to photosystem I in Chlamydomonas reinhardtii. Proc. Natl. Acad. Sci. U. S. A. 115, 3722–3727 (2018).

39. E. H. Harris, D. B. Stern, G. B. Witman, The Chlamydomonas Sourcebook 2nd edn, Academic Press (2009), pp. 1–24.

40. B. D. Engel et al., Native architecture of the Chlamydomonas chloroplast revealed by in situ cryo-electron tomography. eLife 4, 29 (2015).

41. C. Slavov et al., “Super-quenching” state protects Symbiodinium from thermal stress - Implications for coral bleaching. Biochim. Biophys. Acta. 1857, 840–847 (2016).

42. H. J. Liu et al., Phycobilisomes supply excitations to both photosystems in a megacomplex in Cyanobacteria. Science 342, 1104–1107 (2013).

43. S. Akimoto, M. Yokono, E. Yokono, S. Aikawa, A. Kondo, Short-term light adaptation of a cyanobacterium, Synechocystis sp. PCC 6803, probed by time-resolved fluorescence spectroscopy. Plant Physiol. Biochem. 81, 149–154 (2014).

44. C. W. Mullineaux, Function and evolution of grana. Trends Plant Sci. 10, 521–525 (2005).

45. A. R. Grossman, D. Bhaya, K. E. Apt, D. M. Kehoe, Light-harvesting complexes in oxygenic photosynthesis: Diversity, control, and evolution. Annu. Rev. Genet. 29, 231–288 (1995).

46. B. Huang, M. R. Rifkin, D. J. L. Luck, Temperature-sensitive mutations affecting flagellar assembly and function in Chlamydomonas reinhardtii. J. Cell Biol. 72, 67–85 (1977).

47. R. Tokutsu, N. Kato, K. H. Bui, T. Ishikawa, J. Minagawa, Revisiting the supramolecular organization of photosystem II in Chlamydomonas reinhardtii. J. Biol. Chem. 287, 31574–31581 (2012).

48. R. Tokutsu, K. Fujimura-Kamada, T. Yamasaki, K. Okajima, J. Minagawa, UV-A/B radiation rapidly activates photoprotective mechanisms in Chlamydomonas reinhardtii. Plant Physiol. 185, 1894–1902 (2021).

49. R. Tokutsu, M. Iwai, J. Minagawa, CP29, a monomeric light-harvesting complex II protein, is essential for state transitions in Chlamydomonas reinhardtii. J. Biol. Chem. 284, 7777–7782 (2009).

50. R. Tokutsu, K. Fujimura-Kamada, T. Matsuo, T. Yamasaki, J. Minagawa, The CONSTANS flowering complex controls the protective response of photosynthesis in the green alga Chlamydomonas. Nat. Commun. 10 (2019a).

51. R. Tokutsu, J. Minagawa, Energy-dissipative supercomplex of photosystem II associated with LHCSR3 in Chlamydomonas reinhardtii. Proc. Natl. Acad. Sci. U. S. A. 110, 10016–10021 (2013).

