## Supplementary figures and images for "Phosphorylation of light-harvesting complex II controls excitation energy spillover between photosystems"

### Figure S1

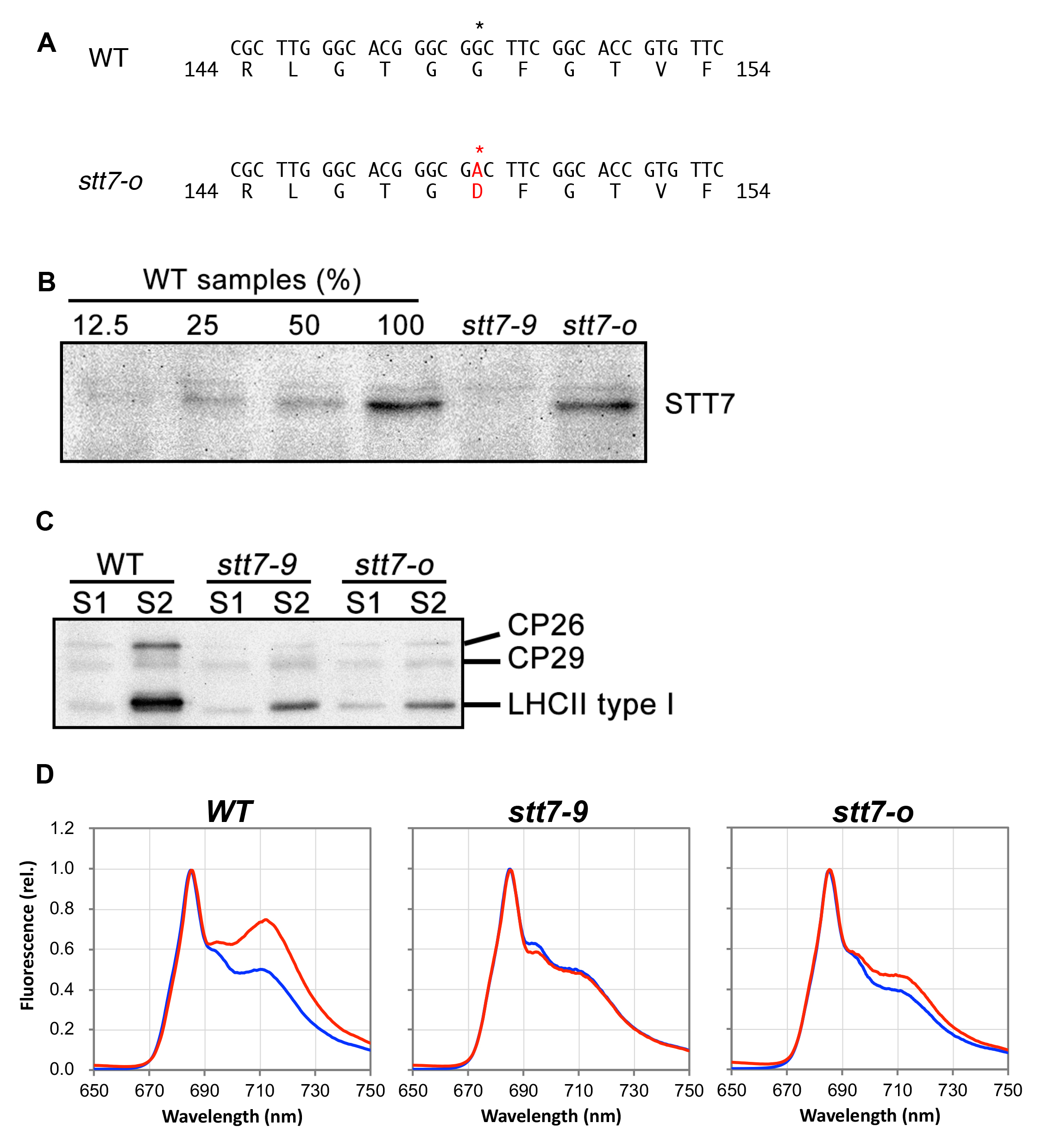

### Figure S2

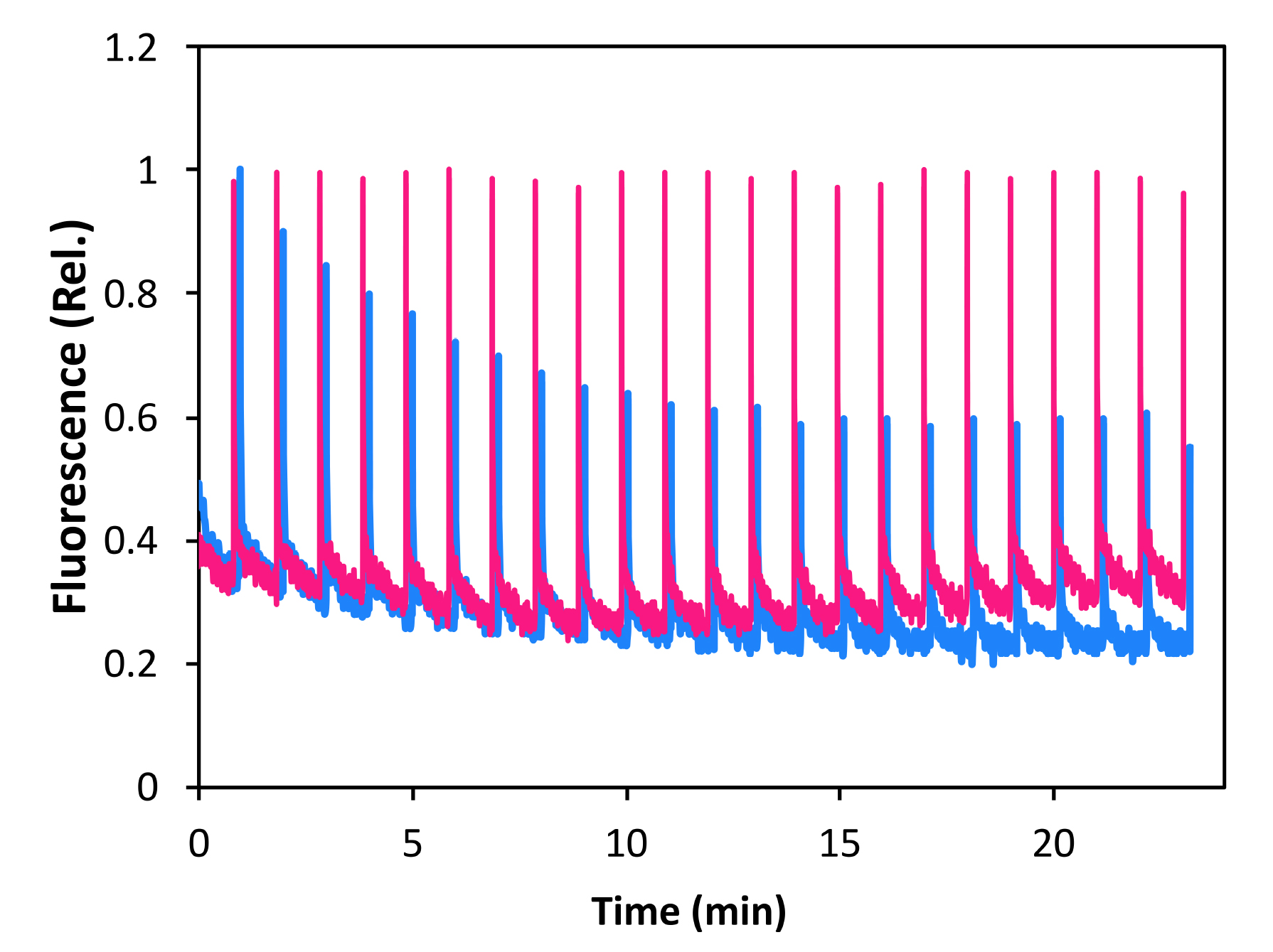

### Figure S3

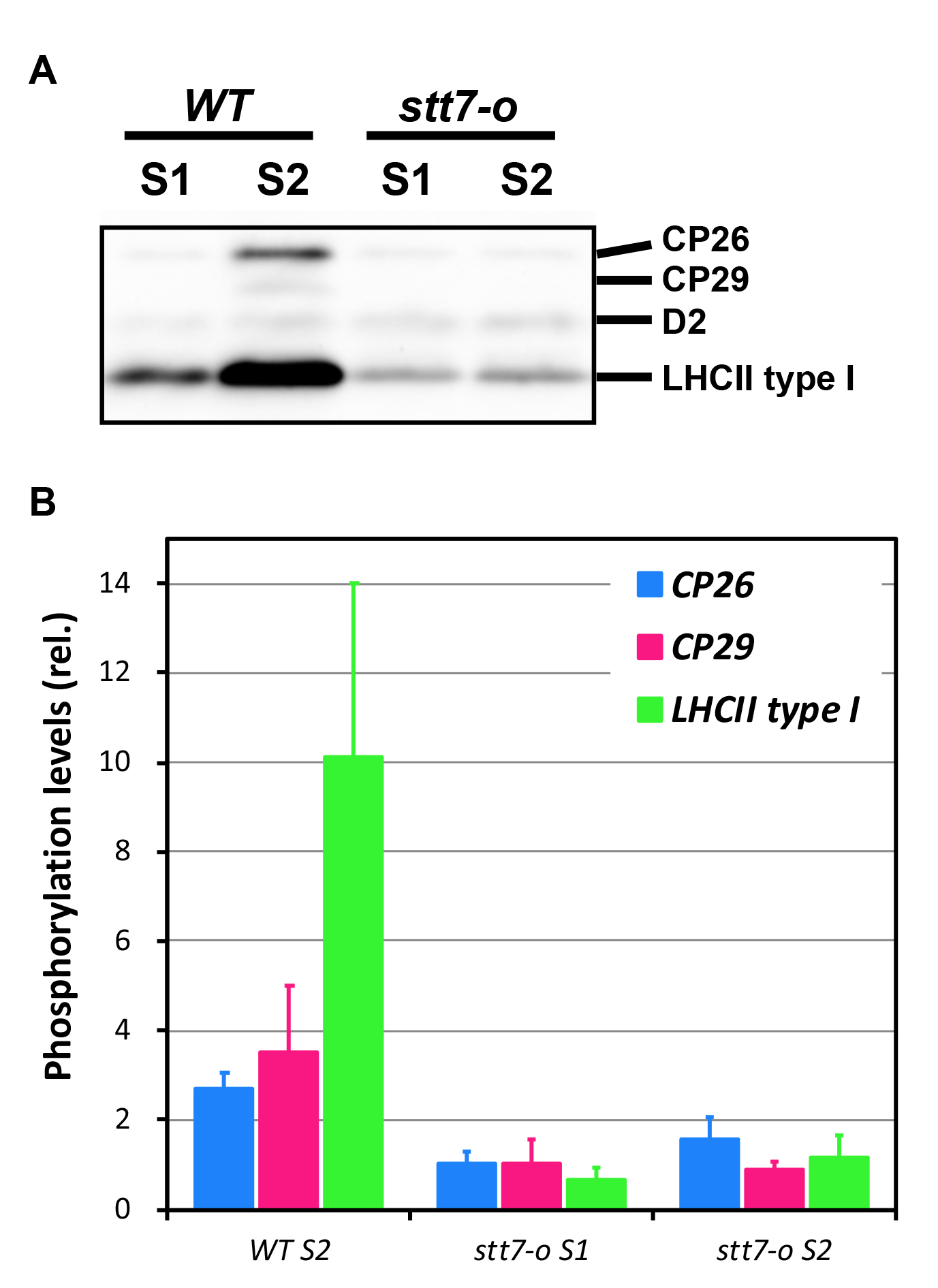

### Figure S4

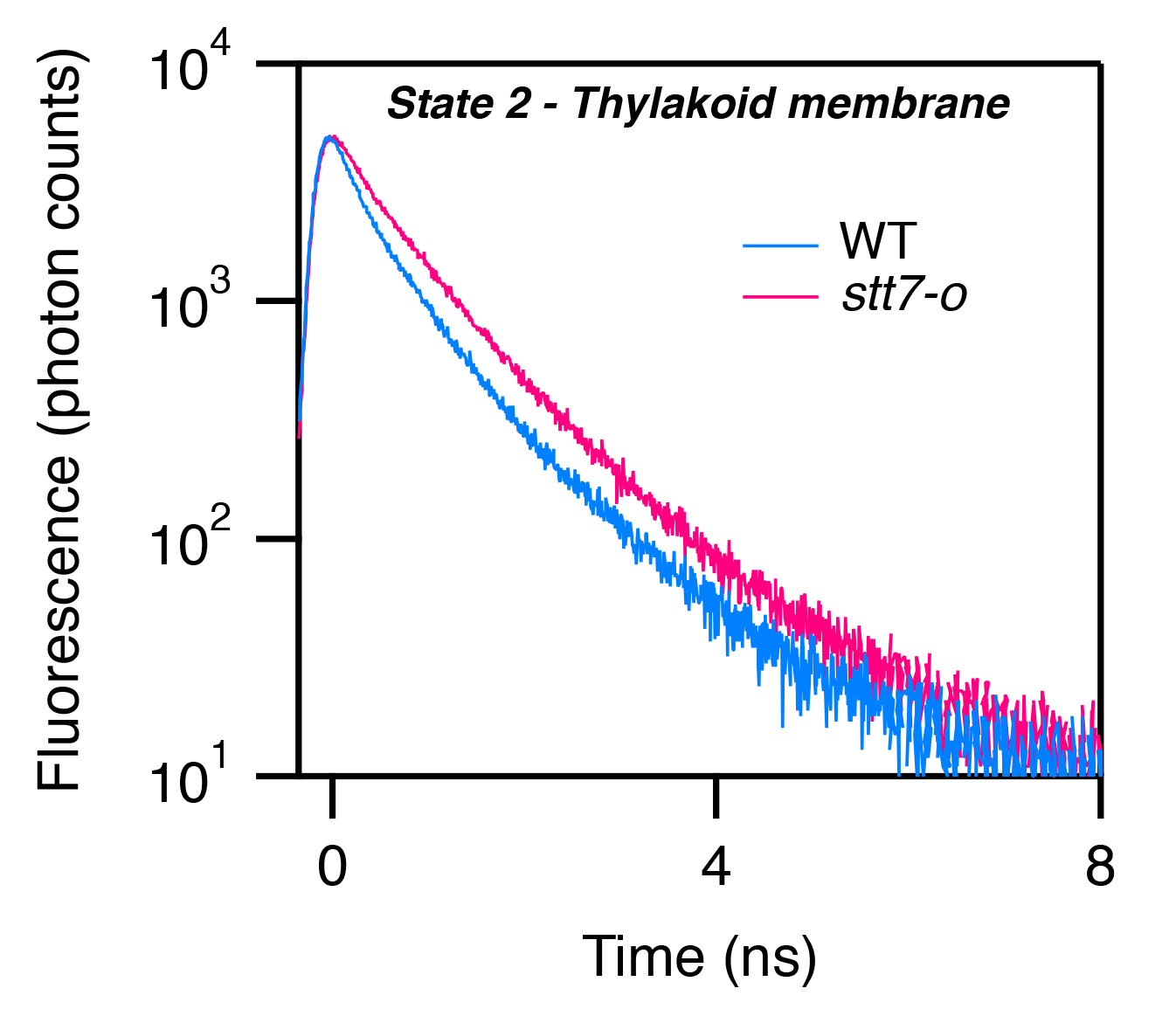

### Figure S5

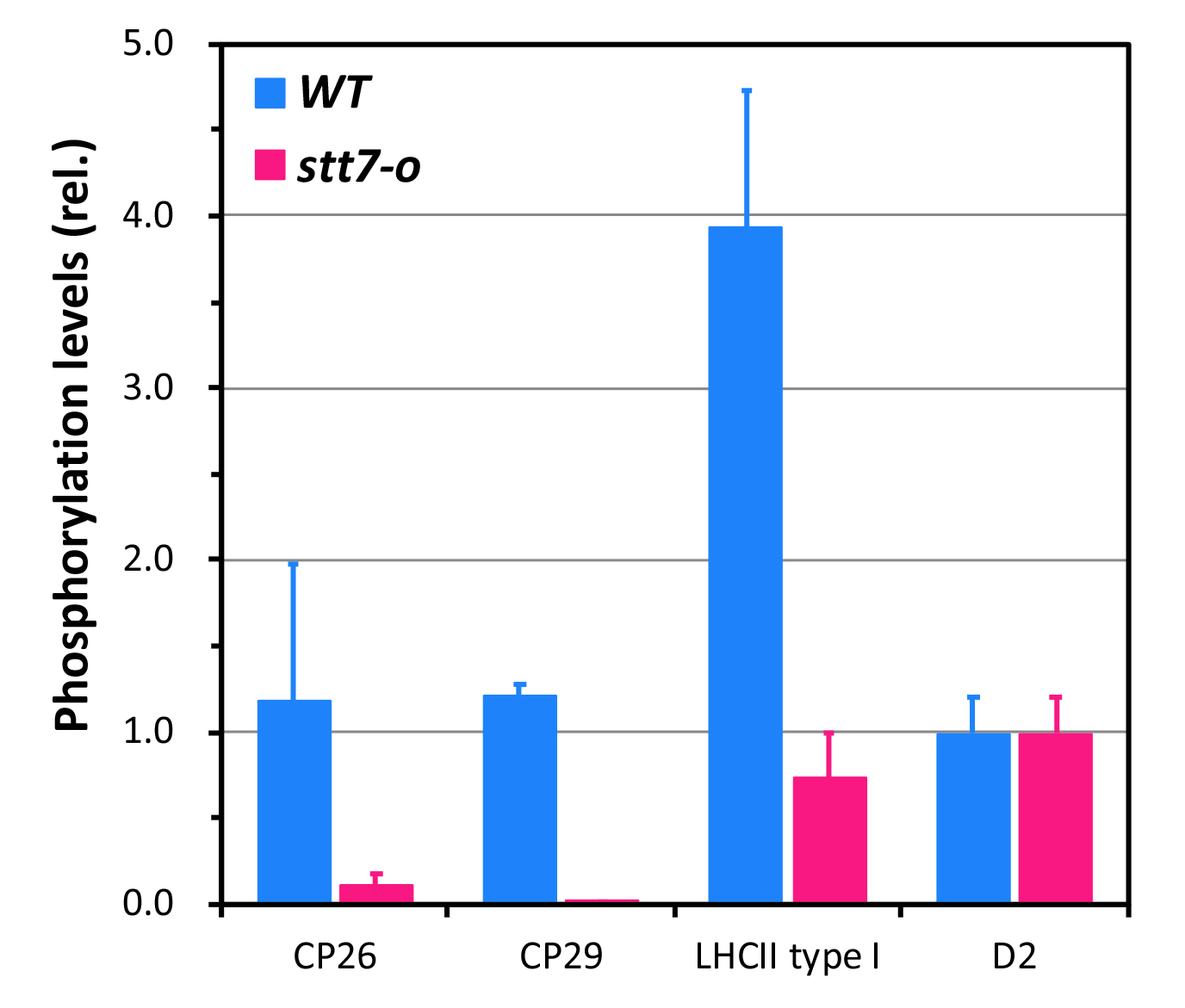

### Figure S6

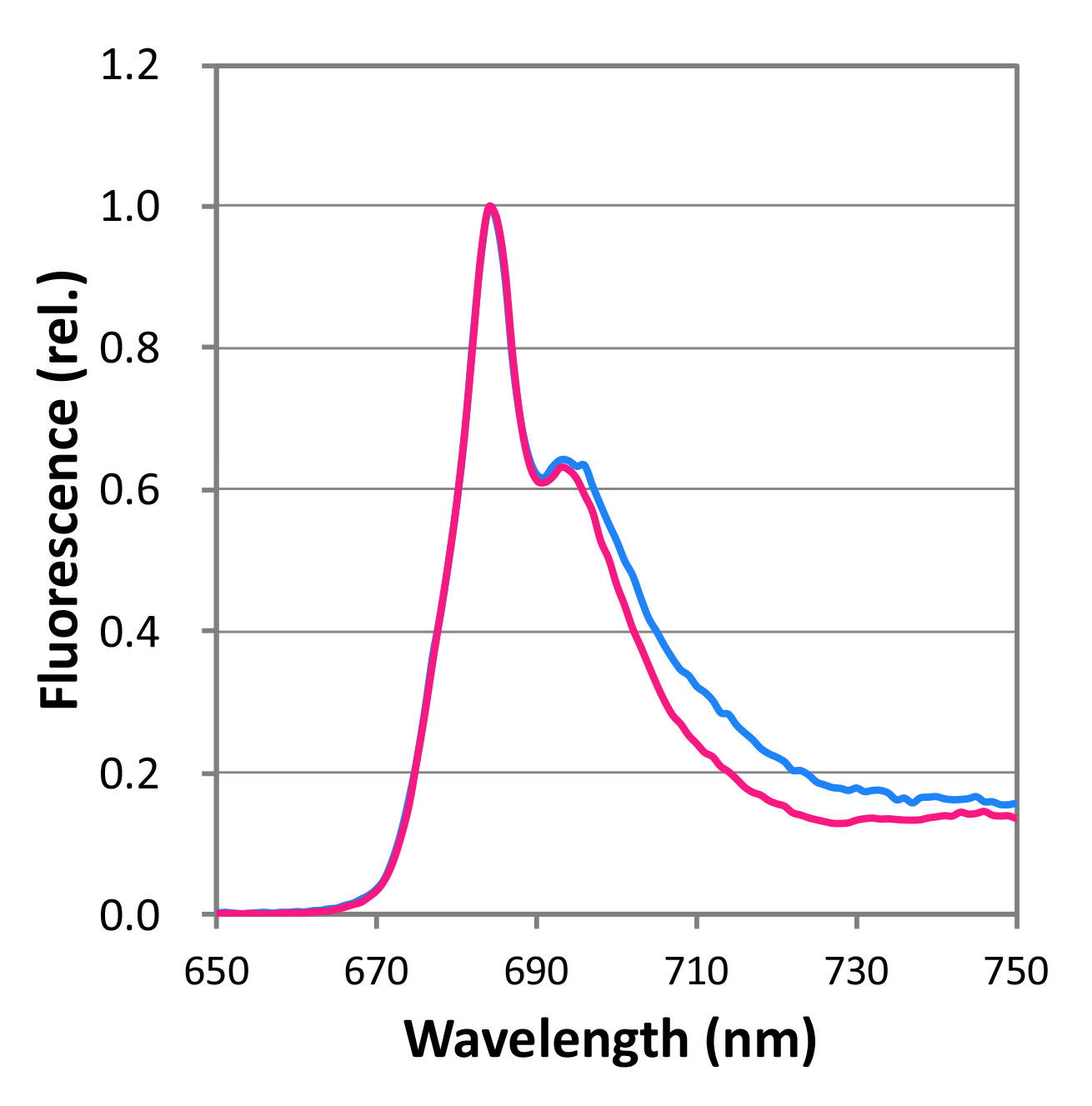

### Figure S7

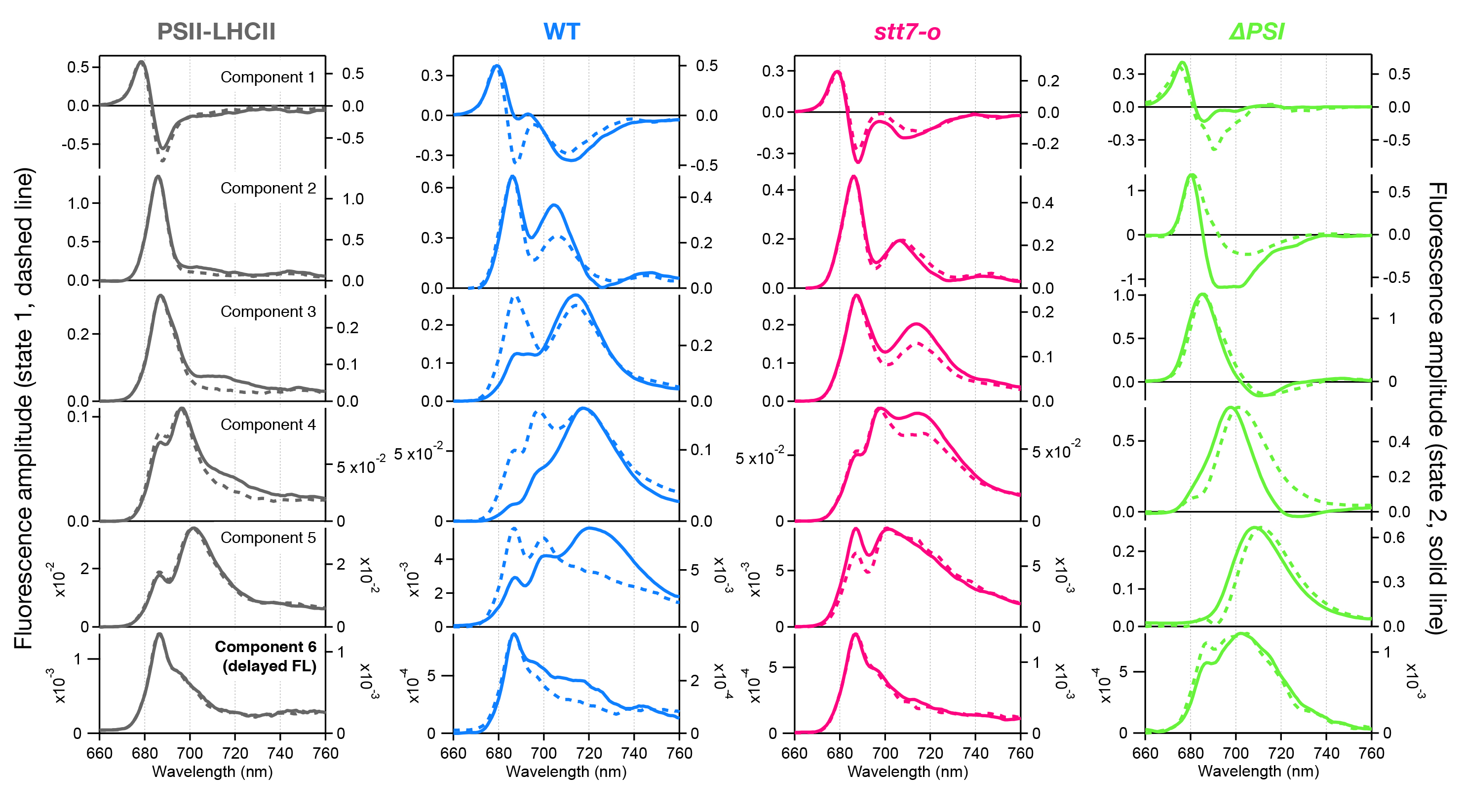

### Figure S8

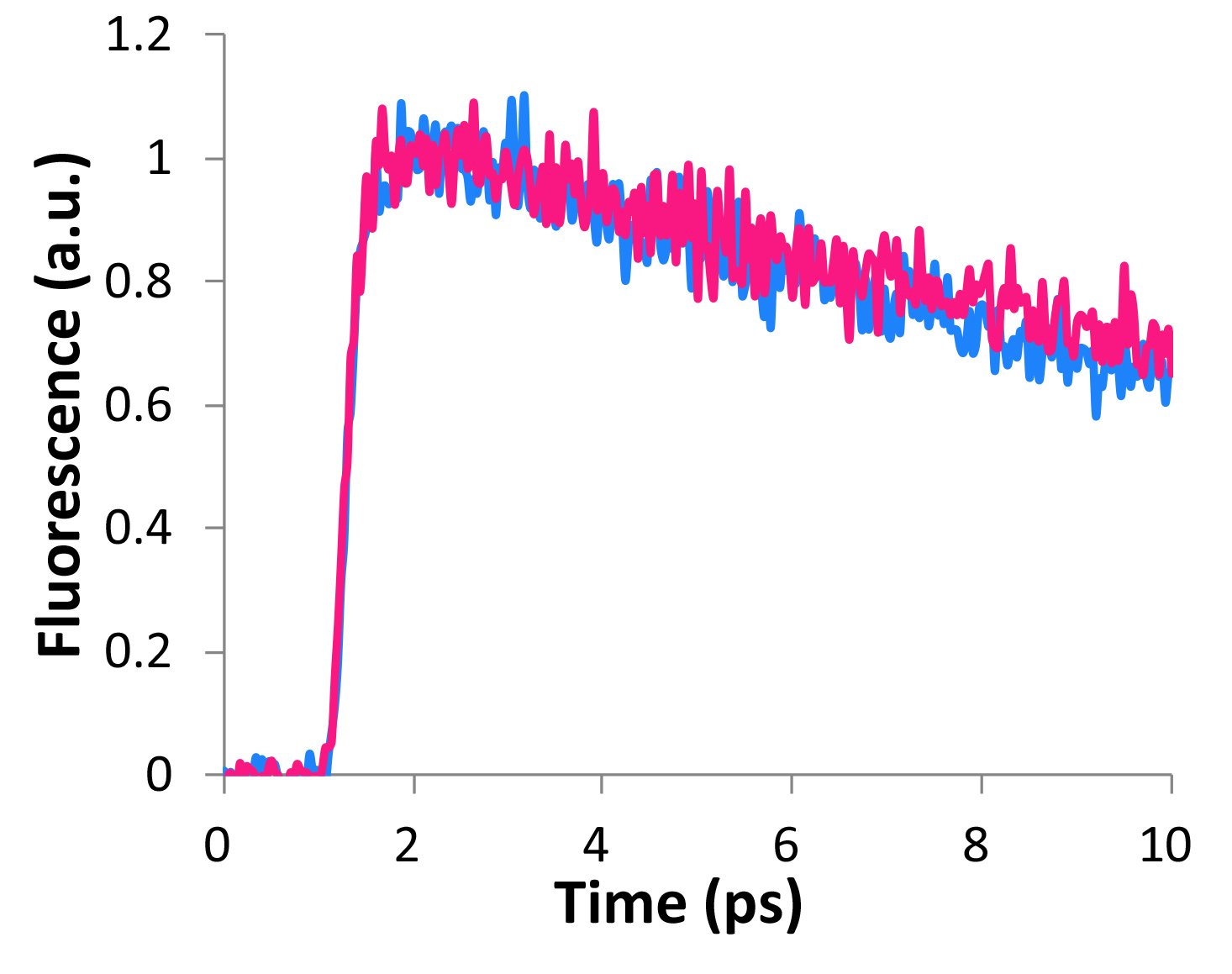

### Figure S9

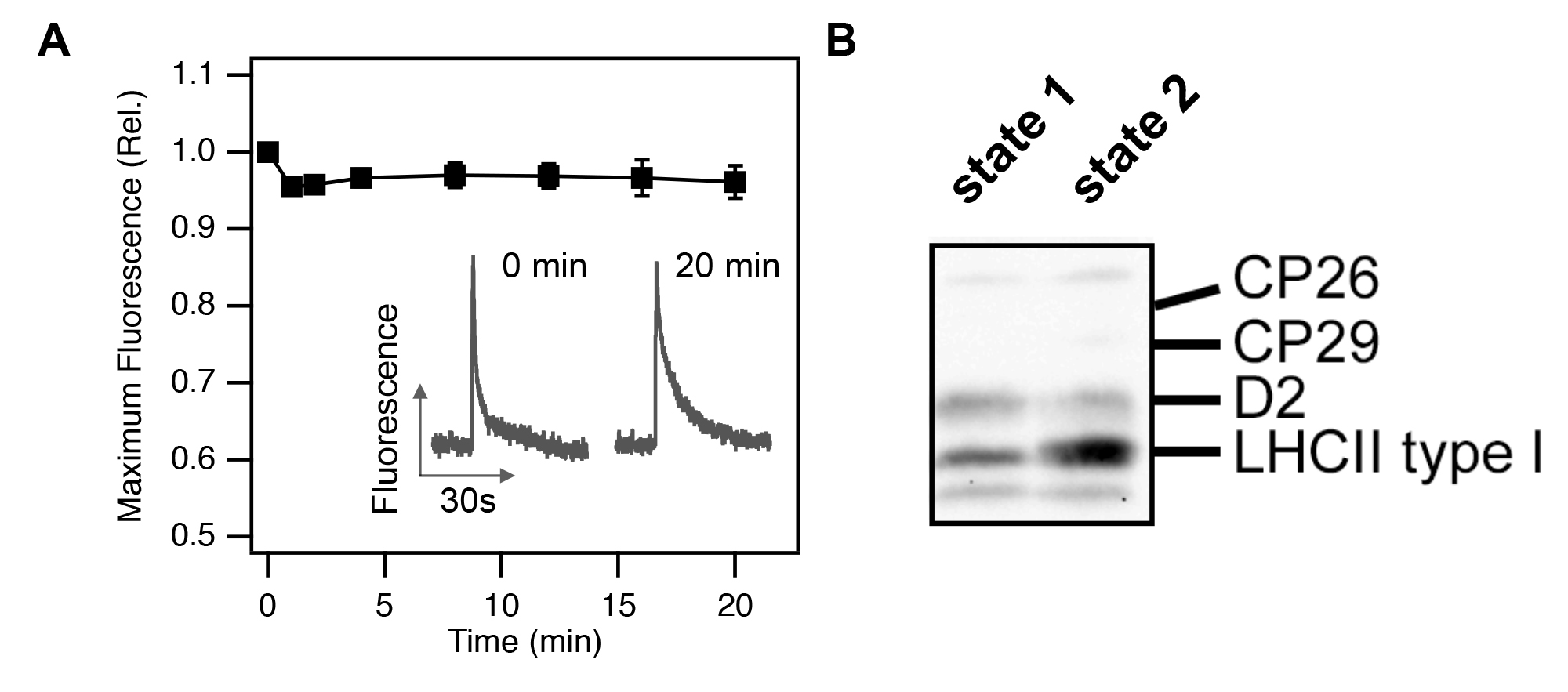

### Table S1

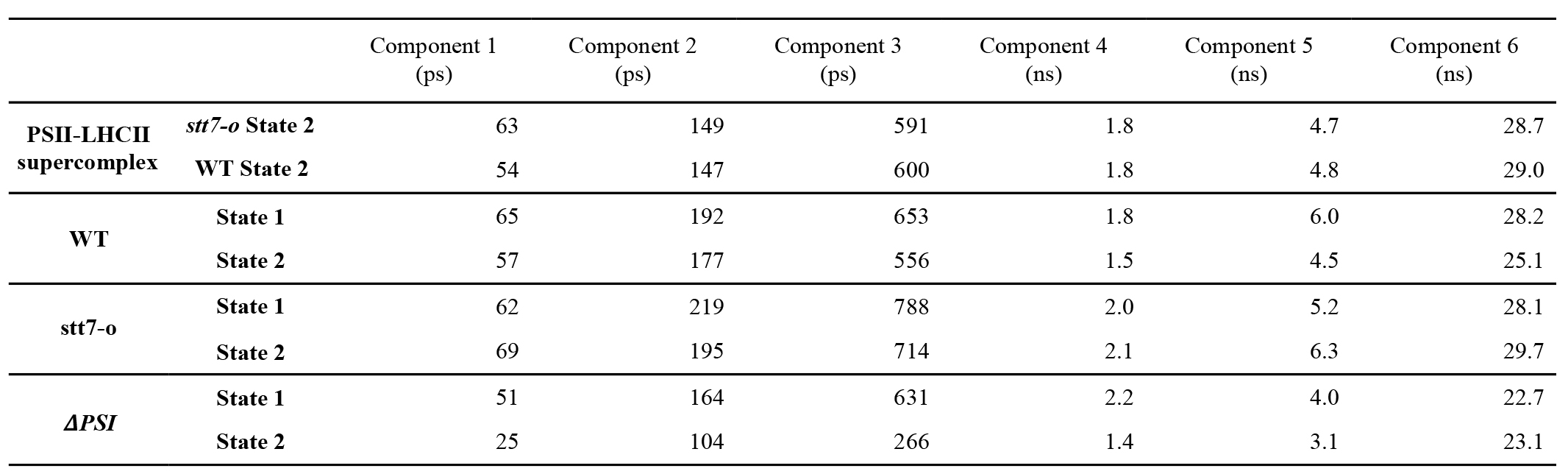

### Table S2

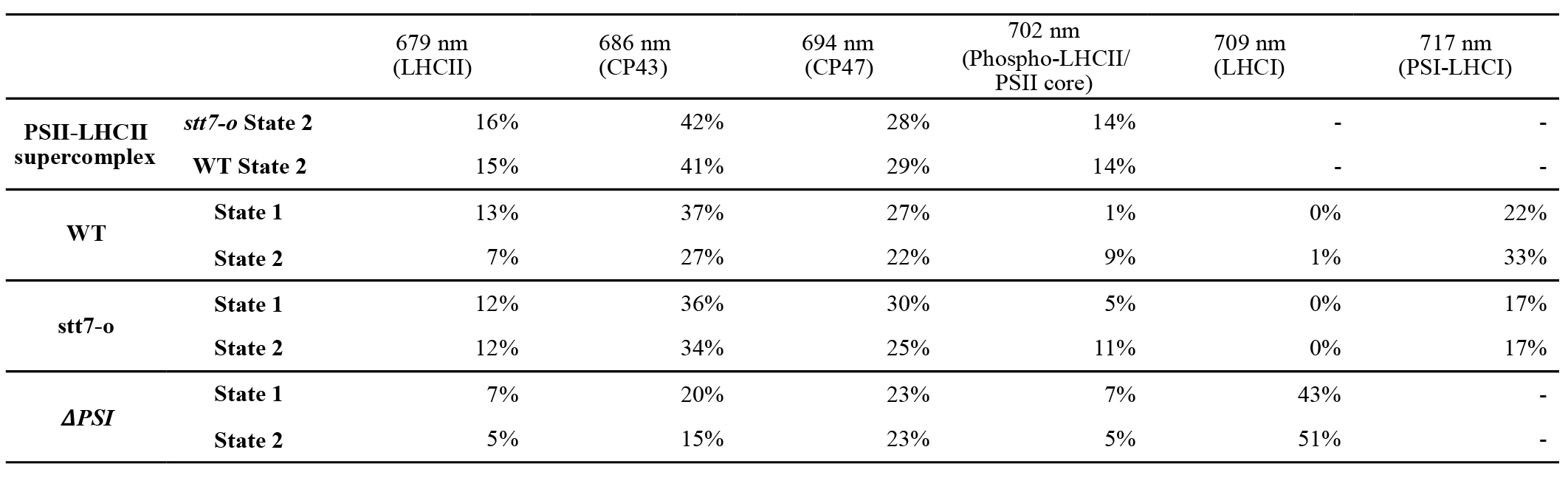
